# Experimental suppression of a keystone protist triggers mesopredator release and biotic homogenization in complex soil microbial communities

**DOI:** 10.1101/2025.04.13.648592

**Authors:** François Maillard, Fredrik Klinghammer, Briana H. Beatty, Hanbang Zou, Enrique Lara, Edith C. Hammer, Anders Tunlid, Peter G. Kennedy

**Author notes:** **Address of corresponding author:** François Maillard.

## Abstract

The keystone species concept suggests that certain members of an ecological community, despite their low abundance, exert disproportionately large effects on species diversity and composition. In microbial ecology, experimental validation of this concept is limited due to significant technical challenges associated with selective species manipulation. Here, we tested this concept within a soil microbial food web by selectively suppressing a protist predator using phototoxicity induced by excessive excitation light during fluorescence microscopy within a microfluidic soil chip system. We targeted a Hypotrichia ciliate taxon—presumed primarily bacterivorous under our experimental conditions—and combined microscopy with metabarcoding of multiple microbial trophic levels to evaluate the effects of this suppression on microbial community abundance, diversity, and composition. Over the 20-day incubation, the chip system supported complex communities of bacteria, fungi, and protists. Following Hypotrichia suppression, two distinct ecological responses were observed: first, an increase in flagellate abundance that was consistent with mesopredator release and accompanied a significant rise in overall protist diversity; second, a convergence in protist community composition, indicative of biotic homogenization. Surprisingly, bacterial community abundance, richness, and composition remained unaffected, likely due to compensatory predation by increased numbers of bacterivorous flagellates. In contrast, fungal diversity decreased following Hypotrichia suppression, presumably resulting from the altered protist communities that favored facultative fungal consumers. Collectively, these findings provide direct experimental evidence that low-abundance microbial predators can function as keystone species, modulating predator community composition and diversity and having cascading effects on lower trophic levels within the brown microbial food web.

## Introduction

Over 50 years ago, Paine introduced the keystone species concept, asserting that certain species, despite their low relative abundance, exert a disproportionate influence on community structure and stability [1, 2]. Since then, this concept has been extensively explored in animal and plant ecology, providing a framework to understand and predict community diversity and composition, with implications for biodiversity and associated ecosystem services [3–5]. In terrestrial and aquatic food webs, apex predators often serve as keystone species, exerting strong top-down control that generates direct and indirect effects across trophic levels, frequently stabilizing entire communities [1–3, 6]. The keystone species concept—defined here as a species whose impact on its community is disproportionately large relative to its abundance [3]—has since expanded from macroorganisms to microorganisms, with applications in microbial ecology [7–10].

In animal microbiome research, notably the human gut microbiome, certain bacterial taxa have been proposed as keystone species because their metabolic or co-metabolic activities drive critical community-level biochemical processes, despite their extremely low abundance [9, 11–13]. The concept has also been applied to environmental microbiomes, particularly in aquatic and terrestrial ecosystems, including soils. Banerjee and colleagues [10] identified up to 200 microbial keystone species from the literature, predominantly bacteria. Despite enthusiasm for the keystone concept in microbial ecology, Röttjers and Faust [14] cautioned that many soil microbial keystone taxa have been identified solely through correlation-based methods—such as DNA metabarcoding network analyses, where highly connected “hubs” are presumed to be keystone species—with limited empirical evidence supporting their keystone roles. Indeed, only 3.5% of proposed microbial keystone species have been experimentally validated [10, 14], contrasting sharply with approximately 75% empirical validation in animal ecology [6]. This discrepancy largely stems from methodological constraints: in animal ecology, manipulative experiments (e.g., selective species removal, often deemed the most robust method for keystone identification [3]) are feasible, whereas selective removal of a single microbial species from complex communities remains challenging.

Interestingly, while animal ecology has typically identified keystone taxa among predator trophic groups relative to their prey [3, 6], soil microbial studies have primarily highlighted primary decomposers—mainly bacteria and, less frequently, fungi—as keystone taxa [10, 15]. This emphasis is surprising, given that soil decomposer food webs, increasingly referred to as “brown food webs” [16], structurally resemble terrestrial macroscopic food webs. These webs encompass multiple trophic levels, with microbial predators, such as protists, preying upon bacterial and fungal primary decomposers [17, 18]. Thus, top-down control by microbial predators could stabilize microbial communities, akin to patterns observed in animal ecology [2, 3]. Given that ciliates are typically the largest and least abundant protist predators in soil communities relative to flagellates and amoebae [19, 20], microbial predator keystone species are likely found among ciliates. Although recent network-based studies have begun identifying potential keystone soil protist taxa, these findings still require experimental validation [21–23].

Here, we experimentally tested the keystone species concept within a microbial brown food web by selectively suppressing a protist predator within a complex microbial community containing other protist predators as well as bacterial and fungal decomposers. We employed microfluidic soil chips designed to mimic natural soil microhabitats by creating synthetic pore spaces interacting with soil. These chips have been shown to be colonized by diverse microbial communities [24, 25], enabling direct microscopic observations and precise, selective elimination of protists using fluorescence microscopy-induced phototoxicity. Our experiments targeted the largest and most morphologically distinct ciliate consistently colonizing the chips, a Hypotrichia species. Hypotrichia exhibit broad feeding ranges, but intermediate-sized species (80–100 µm), like the one targeted in this experiment, primarily prey on bacteria and, to a lesser extent, smaller protists such as flagellates [26]. Thus, we anticipated this Hypotrichia species would predominantly feed on bacteria. Additionally, since ciliates often exhibit preferential feeding on certain bacterial taxa within complex communities [27, 28], we hypothesized that suppressing this Hypotrichia taxon would affect microbial decomposer communities via two trophic pathways. First, removal of the ciliate would release certain bacteria from predation, altering community composition and potentially decreasing bacterial diversity. Second, reduced top-down control would also increase bacterial abundance, intensifying bacterial-fungal competition in favor of bacteria, thereby lowering fungal abundance and diversity. Finally, we hypothesized that suppressing a protist predator would alter the composition and richness of the remaining protist community as other protists filled this ’empty niche,’ significantly shifting community composition and increasing protist diversity. To test these hypotheses, we combined microscopy with DNA-based techniques—including quantitative PCR (qPCR) and high-throughput sequencing (HTS) of bacterial, fungal, and protist taxonomic markers—to assess changes in microbial abundance, diversity, and community composition following ciliate predator suppression.

## Materials and Methods

### Chip design and inoculation

Our microfluidic chips have a cuboid design (14,720 μm × 5,000 μm × 12 μm) with an array of ∼100 μm diameter circular pillars spaced 175 μm center-to-center (see Figure S1a), chosen for rapid inspection at 40× magnification and effective identification and suppression of specific protist taxa. Chips were fabricated following established protocols [25, 29]. Full protocols are available in the Supplementary Materials and Methods. Immediately following PDMS slab bonding—and taking advantage of the temporary hydrophilicity induced by plasma treatment—100 μL of a 4% (mass/volume) fungal necromass suspension sterilized through autoclaving was pipetted directly into the open entrance of each chip. This allowed fungal necromass particles (10 to 30 μm size) to enter the chip via capillary action. Fungal necromass was selected as the primary carbon and nutrient source in the chip because it mimics the organic matter encountered by microorganisms in forest topsoil, representing an ecologically relevant soil microsite where a mycelial network has recently senesced [30]. Moreover, it has been well demonstrated that both bacterial and fungal decomposers participate in the decomposition of fungal necromass, potentially promoting the development of abundant and diverse communities across microbial decomposer domains [31]. We chose *Mortierella alpina* as a necromass source due to the ubiquity of this genus in soils [32]. *M. alpina* was cultured for 28 days in a 250 mL Erlenmeyer flask containing half-strength potato dextrose medium (pH 5). The biomass was harvested, the medium removed and replaced with distilled water and then autoclaved. The autoclaved necromass was washed with distilled water, freeze-dried, and ground into a powder using a mortar and pestle, and subsequently re-suspensed in sterile water, autoclaved, and used to inoculate chips.

After necromass inoculation, the chips were placed in contact with forest topsoil collected from a 60-year-old spruce (*Picea abies*) forest near Lund University’s Stensoffa field station (55.6928° N, 13.4540° E). The soil had a pH of 4.5 and was classified as sandy according to USDA texture classes [33]. Soil was sampled in November 2023 from the A horizon, pooled from five soil cores, and sieved at 2 mm. Approximately 50 g of wet soil was subsampled and stored at 4 °C until use. This forest site was selected because, in a preliminary experiment, we observed consistent colonization by a ciliate species in the subclass Hypotrichia based on morphological characteristics (see below)—from chips inoculated with this soil. Following necromass inoculation in December 2023, 2 g of soil from the pooled sample was placed directly at the chip entrance. The inoculated soil was moistened with sterile-filtered distilled water until apparent saturation (see Figure S1b), and then the Petri dishes sealed with Parafilm and stored in the dark at room temperature for 20 days. The soil in contact with the chip was re-moistened after 10 days with filter-sterilized distilled water. This setup allowed the chip to function as a synthetic, transparent soil pore space system interacting with the soil, thereby facilitating natural microbial movement and colonization over the 20 day incubation period. A total of 20 chips (labeled 1 to 20) were prepared and randomly assigned to one of two groups: a control group and the ciliate suppression group. Additionally, three chips were prepared on coverslips, placed in sealed Petri dishes without soil or necromass, and used as negative controls for potential eDNA contamination during sampling, DNA extraction, PCR, or sequencing.

### Hypotrichia suppression procedure

A protist morphotype, visually identified as belonging to the subclass Hypotrichia, was selected because individuals of this morphotype constituted the largest protists colonizing the chips under our experimental conditions and, due to their low per-chip abundance relative to other protists (around 10 individuals per chip based on preliminary experiments), represented an ideal target to test the keystone concept. The morphology of the observed individuals (elongated, ovoid shaped cells) was also distinct from other ciliates and thus a readily identifiable target. Given the relatively moderate size of the observed Hypotrichia individuals (70 to 110 μm), we anticipated them to feed primarily on bacteria and to a less extent, potentially small flagellates [26]. Under our experimental conditions, Hypotrichia were the largest protists observed, as the ∼80 μm x 10 µm chip entrance (the distance between two pillars times the chip height) restricted the entry of larger ciliates. Moreover, this chip entry sizes mimicked sandy soil pore sizes typically encountered in soil pore systems from the sandy soil selected for this experiment [34].

Over the 20 day incubation, both control and suppression chips (n=10 for each treatment) were inspected daily at 40× magnification to count Hypotrichia individuals. All observations and imaging were performed using an inverted microscope (Nikon Ti2-E inverted microscope with PFS4 hardware autofocus, full 25 mm field-of-view, CoolLED pE300-White MB illumination connected via a 3 mm liquid light guide (LLG), and a Nikon Qi2 camera with 1x F-mount adapter) with 25mm field-of-view. For the suppression treatment, which was applied daily on all Hypotrichia present in the suppression treatment chips, individual Hypotrichia were examined at 400× magnification and exposed to UV range light at excitation of 380-405 nm (CoolLED pE300-White MB illumination) at 100% intensity for 5 seconds. This exposure triggered cell death—likely by reacting with photoreceptive compounds triggering oxidative stress and compromising membrane integrity (Figure 1a). Hypotrichia individuals were monitored by bright-field microscopy until cell breakdown was evident. If Hypotrichia death did not occur after the initial exposure, an additional 2-second exposure was applied, and the procedure repeated until suppression was observed. Notably, photo-exposed Hypotrichia individuals formed vacuoles, indicating that their cytoplasmic content was not fully released into the chip, thereby minimizing potential bias from providing additional food sources for bacteria and fungi due to protist cell death (Figure 1a). Five chips were excluded from subsequent microscopy analyses due to technical issues: one control chip (chip 18) exhibited a coverslip fissure on day 20 and was used only for eDNA-based techniques; one suppression chip (chip 19) was omitted after PDMS unbonding was observed in the top left corner; and three additional chips (chips 4 and 5 from the suppression group, and chip 13 from the control group) were excluded due to very low Hypotrichia colonization (see Table S1 for a summary of the chips retained for analysis).

**Figure 1.**
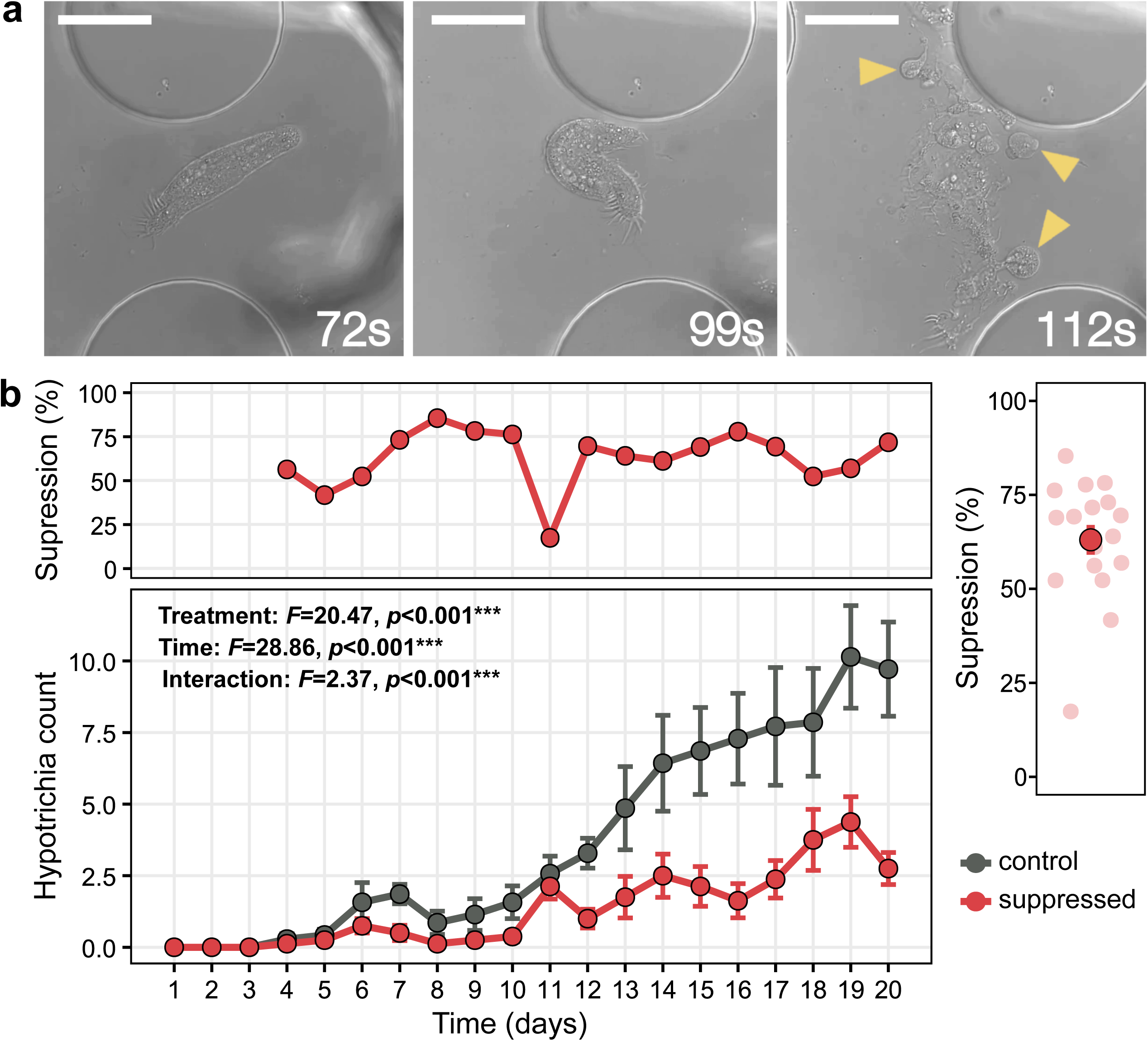
Effects of the suppression procedure on Hypotrichia abundance. (a) Representative microscopy images showing Hypotrichia in chips inoculated with forest topsoil at 72, 99, and 112 seconds following a 5-second exposure to 395 nm excitation light (CoolLED pE300-White MB illumination). Yellow arrows indicate post-senescence vacuole formation. Scale bar = 50 µm. (b) Bottom panel shows daily Hypotrichia individual counts per chip over 20 days (averaged by treatment: control vs. suppressed). In the suppressed treatment (red line), all the counted Hypotrichia have been suppressed. The effects of treatment, time (days post-inoculation), and their interaction were evaluated using a linear mixed-effects model (time as categorical, chip as random factor) via Type III ANOVA with Satterthwaite’s method (\**p* < 0.05, \*\**p* < 0.01, \*\*\**p* < 0.001). The top panel shows average Hypotrichia suppression percentage over time, and the right panel shows average suppression percentage across all time points. Data are mean ± SE.

### Microscopy and image analysis of the chips

At 20 days post-inoculation, we recorded 15 second videos at 400× magnification to count bacteria, fungi, and protists with real-time videos recorded via a digital camera (USB29 UXG M). Video capture, rather than images, enabled more accurate identification of protists based on their characteristic movements, which helped distinguish them from soil debris or fungal necromass. For each chip, 100 videos of 15 seconds each were recorded from open spaces between pillars. Specifically, 20 videos were recorded per row at intervals corresponding to every 5 pillars from the entrance, ensuring a representative coverage of the chip. Videos were recorded and assigned random numbers by F.M., and subsequent counting of bacteria, fungi, and protists was performed by F.K. under a single-blind setup to reduce observational bias. For bacterial abundance, a scoring system ranging from 1 (very low abundance) to 5 (very high abundance) was used (see Figure S2 for representative images). In addition to the scoring system for bacterial abundance, we also utilized a newly developed deep-learning algorithm as previously described [24]. As the deep-learning algorithm cannot well quantify bacterial clusters and dividing bacteria, to avoid small necromass pieces interfering with quantifications, only individual bacterial cells were retained. Moreover, we applied an inclusivity cut-off on bacterial cell maximum length from 0.3 to 5 µm [24]. The bacterial density (number of cells per frame) was then summed based on the 100 images per chip (see Figure S3 for a representative example of a processed image).

Fungal hyphae were counted as single units when observed crossing the frame independently, to avoid multiple counts of branching hyphae. Protists were counted and categorized into functional groups: ciliates (moving by cilia), amoebae (exhibiting amoeboid movement), and flagellates (moving via flagella; amoeboflagellates like cercomonads were grouped with flagellates) (Figure S4). Although this functional group classification does not necessarily reflect evolutionary relationships (except for ciliates), it provides a basis for inferring functional roles (e.g., flagellates as primarily bacterial feeders, amoebae as more omnivorous), and has been widely used in literature allowing direct comparisons with published research [19, 20].

### Chip eDNA extraction, and high-throughput amplicon sequencing and microbial qPCR

After video recording at day 20, residual soil was removed, and the chip along with its coverslip was cleaned to reduce eDNA contamination. The chip was then detached and processed for genomic DNA extraction using the PowerSoil Pro Kit (Qiagen, Hilden, Germany) following the manufacturer’s protocol (full details provided in the Supplementary Materials). Bacterial abundances were quantified by qPCR using the 16S rRNA gene (primers 1401F/968R [35]) on a Stratagene Mx3005P system, although fungal qPCR (primers FR1/FF390 [36]) was unsuccessful due to low genomic yields. Microbial community structure was characterized by high-throughput sequencing of taxonomic markers from bacteria (16S V4 region using 515F–806R [37]), fungi (ITS2 region using 5.8S-Fun/ITS4-Fun [38]), and protists (18S rRNA gene using 616*f–1132r [39]) from 16 chip samples and three non-inoculated chip controls. Sequencing reads were processed using the DADA2 pipeline in R [40]—including quality filtering, chimera removal, and OTU clustering at 97% similarity—and taxonomically assigned using the SILVA [41], UNITE [42], and PR2 [43] databases, with additional filtering to remove contaminants and non-target sequences. To examine the effects of Hypotrichia suppression on protist community composition and diversity, all Hypotrichia sequences were removed from the protist dataset before analyses, a summary of the chips retained is provided in Table S1, and protist OTUs were regrouped into functional groups based on taxonomic information (see Table S2). Full protocols are available in the Supplementary Materials and Methods.

### Data analyses

All statistical analyses and data visualizations were performed in R [44]with a significance threshold of α = 0.05. To assess the effect of the suppression procedure on Hypotrichia abundance over time, linear mixed-effects models (using the lme4 package [45]) were employed with “chip” as a random factor to account for repeated measures. Fixed effects were evaluated using Type III ANOVA with Satterthwaite’s method (lmerTest package [46]). OTU richness, Simpson’s diversity, and Simpson’s evenness for bacterial, fungal, and protist communities were calculated using the Vegan package [47]. For comparisons of microbial abundance and diversity metrics, normality was first tested using the Shapiro–Wilk test; non-normal variables were log-transformed. Homoscedasticity was then assessed using Bartlett’s test; if variances were homogeneous, a t-test was used, otherwise a Welch’s t-test was applied. The effects of Hypotrichia suppression on community composition were analyzed using permutational multivariate analysis of variance (PERMANOVA; Vegan package) based on Bray–Curtis dissimilarities, with results visualized via non-metric multidimensional scaling (NMDS). Pearson’s correlation was used to assess relationships between bacterial, fungal, and protist diversity and Hypotrichia counts at day 20 post-inoculation, as well as between the abundance (microscopy) and relative abundance (metabarcoding) of flagellates, amoebae, and non-Hypotrichia ciliates and Hypotrichia counts.

## Results

Microscopy at 40× magnification revealed that Hypotrichia individuals began colonizing chips by day 4 post-inoculation (Figure 1b). In control chips, their numbers increased significantly over time (*F* = 28.86, *p* < 0.001), reaching peak abundances by day 19 (Figure 1b). The suppression treatment significantly reduced Hypotrichia abundance (*F* = 20.47, *p* < 0.001), with suppression efficiencies ranging from 23% to 75%, averaging 63% throughout the experiment. Metabarcoding of protist communities at day 20 showed a 40% reduction in Hypotrichia relative abundance in the suppressed treatment compared to controls, though this difference was not statistically significant (Figure S5a). Six distinct Hypotrichia OTUs were identified, with three pairs exhibiting high sequence similarity (93–95%; Figure S5b). Searches against the PR2 database matched these OTUs to either Oxytrichidae or Pseudourostylidae. Notably, OTU30 was dominant, accounting for 74% of Hypotrichia sequences and appearing in 69% of chips, whereas most other Hypotrichia OTUs were detected in only one chip (Figure S5c).

Bacterial abundance, assessed via deep-learning analysis of microscopy videos (400× magnification, 100 videos per chip) at day 20 post-inoculation, increased by an average of 80% in Hypotrichia-suppressed chips, although this was not statistically significant (Figure 2a). This trend was confirmed by human-based visual scoring (Figure S7) and bacterial qPCR (Figure 2b). In contrast, bacterial OTU richness (Figure 2c), Simpson’s diversity (Figure 2d), and evenness (Figure 2e) were largely equivalent between treatments. NMDS analysis of bacterial community composition revealed no clear treatment-related clustering, a result supported by the non-significant PERMANOVA test (Figure 2f). Gammaproteobacteria dominated bacterial communities (up to 70% of sequences), followed by Alphaproteobacteria, Actinobacteria, and Bacteroidia, with no marked class-level differences between treatments (Figure 2g). The dominant bacterial genera included *Cavicella*, *Alkanindiges* (both Gammaproteobacteria), *Burkholderia* (Betaproteobacteria), *Mycobacterium* (Actinobacteria), *Roseateles* (Betaproteobacteria), *Chitinophaga*, *Mucilaginibacter* (both Bacteroidota), and *Herminiimonas* (Betaproteobacteria) (Figure S8).

**Figure 2.**
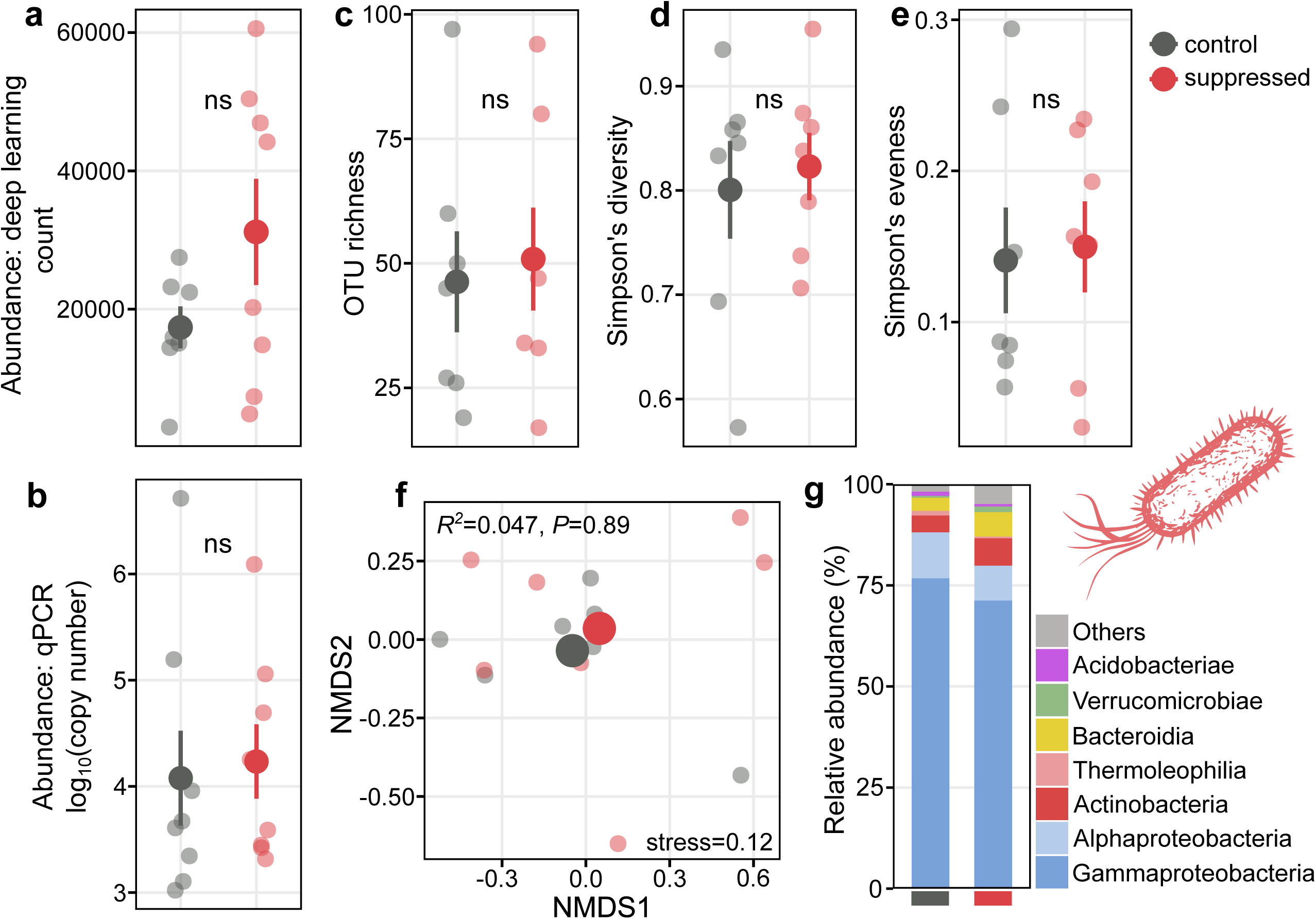
Effects of Hypotrichia suppression on bacterial communities. (a) Bacterial abundance based on microscopy combined with a deep-learning bacterial cell identification and counting algorithm (summed from 100 images per chip), (b) bacterial abundance based on 16S qPCR per chip, (c) OTU richness, (d) Simpson’s diversity, and (e) Simpson’s evenness from 16S metabarcoding. Large dark dots indicate means ± SE; lighter dots represent individual chips. Statistical comparisons made using t-tests or Welch’s tests as appropriate. (f) Bacterial community composition (rarefied OTU abundance) shown by non-metric multidimensional scaling (NMDS) using Bray–Curtis dissimilarity. Large dots are centroids; small dots individual communities. PERMANOVA tested treatment effects (\**p* < 0.05, \*\**p* < 0.01, \*\*\**p* < 0.001). (g) Relative abundance of the seven most abundant bacterial classes averaged by treatment.

Fungal abundance, assessed at the day 20 through microscopy (400× magnification, summed from 100 videos per chip), was marginally lower in the Hypotrichia suppression treatment compared to controls (*p* = 0.07; Figure 3a). Metabarcoding analyses also indicated marginally reduced fungal OTU richness (*p* = 0.07) and a significantly lower Simpson’s diversity in the suppressed treatment (*p* = 0.04; Figure 3b,c). NMDS analysis of fungal OTU composition indicated some separation of treatment centroids, though the PERMANOVA did not find a significant difference (Figure 3e). The fungal community included Ascomycota (Eurotiomycetes and Leotiomycetes), Basidiomycota (Tremellomycetes yeasts), and early-diverging lineages (Mortierellomycetes and Rozellomycota; Figure 3f). The dominant fungal genera encompassed both filamentous and yeast forms, including *Aspergillus* and *Penicillium* (both Eurotiomycetes), *Oidiodendron* (Leotiomycetes), Solicoccozyma, Saitozyma (both Tremellomycetes), *Mortierella* (Mortierellomycota), and *Nadsonia* (Saccharomycetes; Figure S9).

**Figure 3.**
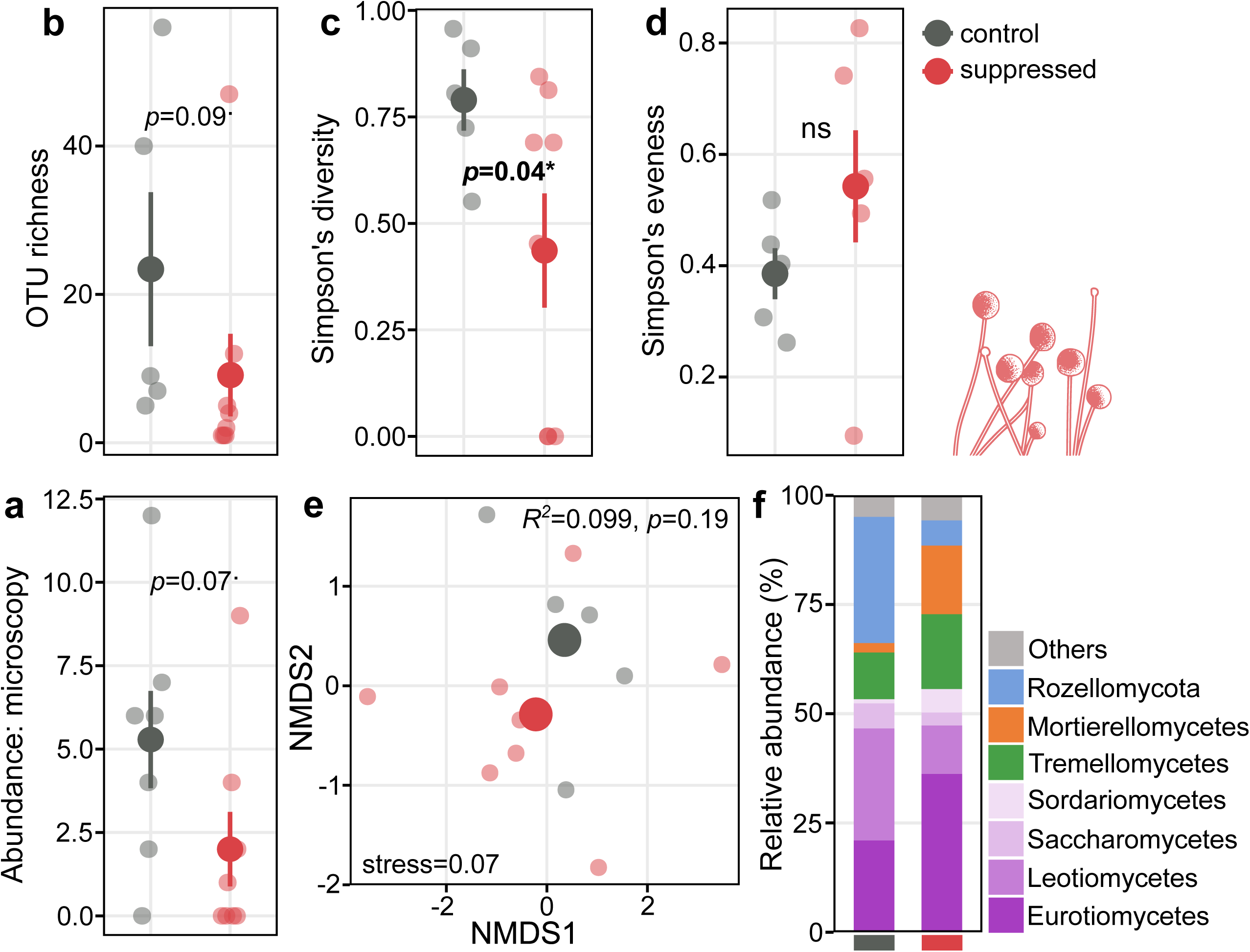
Effects of Hypotrichia suppression on fungal communities. (a) Fungal abundance from microscopy (summed from 100 images per chip), (b) OTU richness, (c) Simpson’s diversity, and (d) Simpson’s evenness based on ITS metabarcoding. Large dark dots indicate means ± SE; lighter dots represent individual chips. Statistical comparisons were made using t-tests or Welch’s tests as appropriate. (e) NMDS analysis (Bray–Curtis dissimilarity) of fungal OTU composition (large dots are centroids; small dots individual chips). Treatment effects compared by PERMANOVA (\**p* < 0.05, \*\**p* < 0.01, \*\*\**p* < 0.001). (g) Relative abundance of the seven most abundant fungal classes averaged by treatment.

Protist OTU richness at day 20, assessed via metabarcoding, did not differ significantly between treatments (Figure 4a). However, both Simpson’s diversity (*p* = 0.007; Figure 4b) and evenness (*p* = 0.045; Figure 4c) were significantly higher in the suppression treatment. PERMANOVA also indicated a significant effect of Hypotrichia suppression on protist OTU composition (*p* = 0.04). Specifically, the NMDS analysis showed higher variability in protist communities of control chips compared to suppressed chips, which had more homogeneous community compositions (Figure 4d). Treating Hypotrichia abundance as a continuous variable (40× microscopy counts at day 20) confirmed this pattern, with decreasing Hypotrichia counts correlating with increased similarity in protist community composition among chips (Figure 4e). The control chips were dominated by Amoebozoa, Heterolobosea, Ciliophora, and Cercozoa, while suppressed chips contained substantial abundances of Euglenozoa, Bigyra, and Choanoflagellata (Figure 4f). Additional microscopic eukaryotes were also detected via metabarcoding. Rotifera occurred at very low relative abundances and only in a single sample (Figure S6), while nematode sequences represented approximately 40% of the total 18S sequences, with no significant differences between treatments (Figure S6).

**Figure 4.**
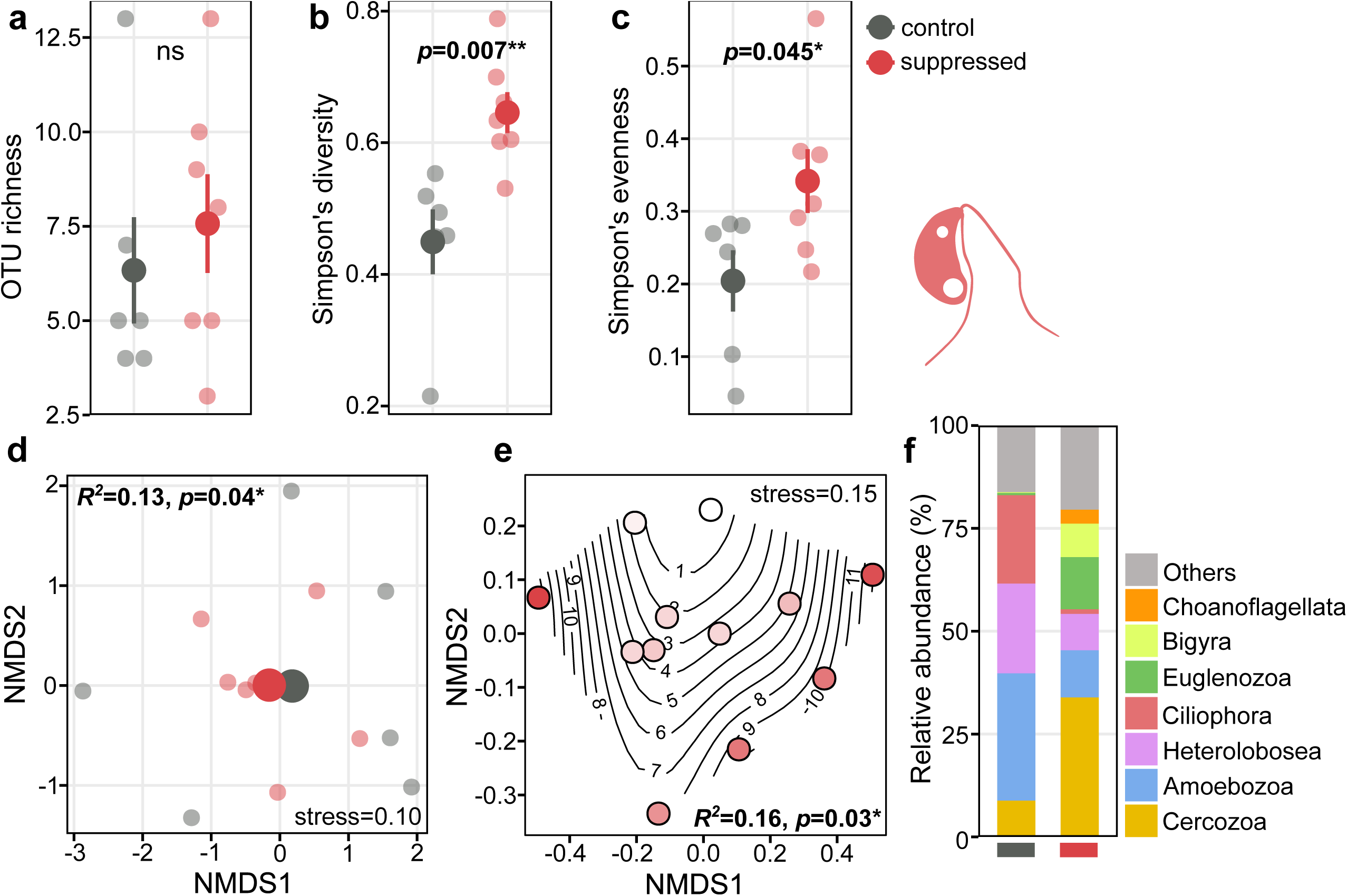
Effects of Hypotrichia suppression on protist communities assessed via 18S metabarcoding. (a) OTU richness, (b) Simpson’s diversity, and (c) Simpson’s evenness (large dots means ± SE; small dots individual chips; statistical comparisons by t-tests or Welch’s tests). (d) Protist community composition (NMDS with Bray–Curtis dissimilarity; large dots centroids, small dots individual communities). (e) NMDS with Hypotrichia counts at day 20 as a continuous variable (circles colored by Hypotrichia abundance, darker indicates higher abundance; segments indicate specific counts per chip). Treatment effects tested via PERMANOVA (\**p* < 0.05, \*\**p* < 0.01, \*\*\**p* < 0.001). (g) Relative abundance of the seven most abundant protist phyla/classes averaged per treatment.

Microscopy-based counts (400× magnification, summed from 100 videos per chip) indicated an average protist abundance of ∼280 individuals in controls versus ∼400 in suppressed treatments, though this difference was not statistically significant (Figure 5a). Amoebae and flagellates dominated both microscopy (Figure 5a) and metabarcoding datasets (Figure 5b). Flagellate abundance did increase significantly with Hypotrichia suppression (microscopy: Figure 5c; metabarcoding: *p* = 0.001, Figure 5d), while amoebae abundance was not significantly affected by suppression in either analysis (microscopy: Figure 5e; metabarcoding: Figure 5f). Conversely, non- Hypotrichia ciliates significantly decreased in microscopy analyses (*p* = 0.02), though the trend was non-significant in metabarcoding data. At finer taxonomic resolution, the dominant flagellate taxa (e.g., MAST-12C, *Neobodo*, *Monosiga*, *Cercomonas*, *Bodo*, and *Glissomonadida*) increased with suppression, whereas dominant non-Hypotrichia ciliates (*Platyophrya*, *Microthoracida*) decreased. Amoebae taxa exhibited variable responses across taxa (Figure S10).

**Figure 5.**
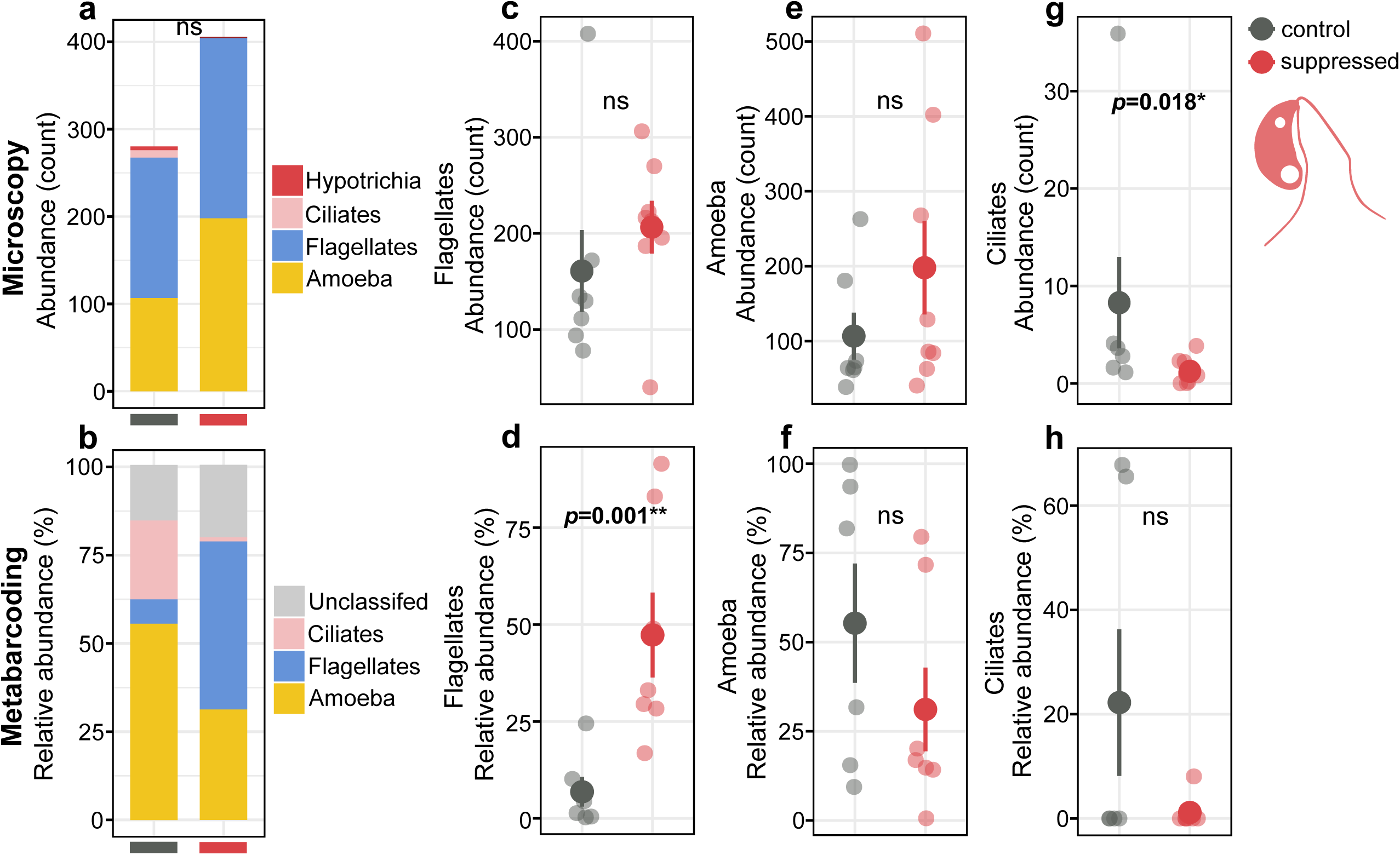
Microscopy-based effects of Hypotrichia suppression on protist abundance: (a) total protist count (summed from 100 images per chip; Hypotrichia not included in “ciliates”), (c) flagellate abundance, (e) amoeba abundance, and (g) ciliate abundance. Metabarcoding-based effects of Hypotrichia suppression on (b) relative abundance of OTUs and specifically for (d) flagellates, (f) amoebae, and (h) ciliates. Statistical comparisons performed via t-tests or Welch’s tests (\**p* < 0.05, \*\**p* < 0.01, \*\*\**p* < 0.001).

When analyzed as a continuous variable, Hypotrichia abundance at day 20 showed no significant relationship with bacterial (Figure 11a) or fungal Simpson’s diversity (Figure 11b), but was significantly negatively correlated with protist Simpson’s diversity (*p* = 0.03; Figure 11c). Hypotrichia abundance was also significantly negatively correlated with flagellate abundance in both the microscopy (*p* < 0.01; Figure 6a) and metabarcoding (*p* < 0.01; Figure 6b) analyses. Amoebae showed no significant correlation in either analysis (microscopy: Figure 6c; metabarcoding: Figure 6d). Finally, non-Hypotrichia ciliates were significantly positively correlated with Hypotrichia abundance in the microscopy analysis (*p* = 0.03; Figure 6e), but not in the metabarcoding analysis (Figure 6f).

**Figure 6.**
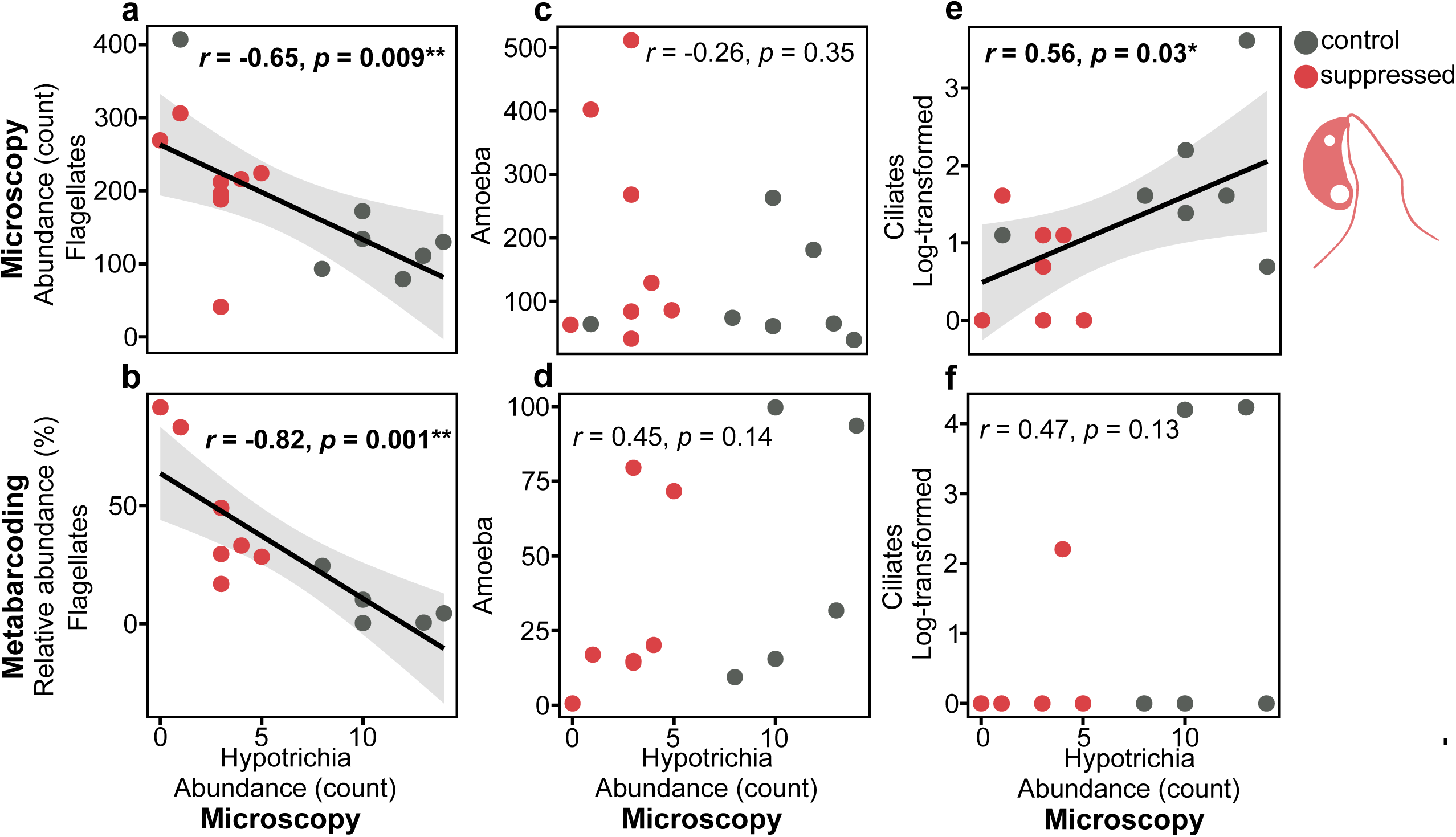
Pearson’s correlations between Simpson’s diversity indices of (a) bacterial, (b) fungal, and (c) protist communities and Hypotrichia abundance (day 20, microscopy-based). Asterisks denote significance (\**p* < 0.05, \*\**p* < 0.01, \*\*\**p* < 0.001)

## Discussion

Here, we used soil chips mimicking a soil environment with micrometer-sized pore spaces to support the establishment and growth of complex microbial communities—including bacteria, fungi, and protists—in semi-controlled conditions, thereby enabling targeted suppression of a protist predator via phototoxicity. To our knowledge, this experiment represents the first experimental attempt to determine the potential keystone role of a microbial predator species through selective suppression.

Complex interacting microbial communities were successfully established, with average species richness of 48 bacterial OTUs, 14 fungal OTUs, and 7 protist OTUs. These communities spanned broad phylogenetic diversity—including Proteobacteria, Actinobacteria, and Bacteroidota among bacteria; Eurotiomycetes, Leotiomycetes, Tremellomycetes, and Mortierellomycetes among fungi; and Cercozoa, Amoebozoa, Heterolobosea, and Ciliophora among protists—all abundant lineages within soil microbial communities [31, 48–50]. Moreover, relative abundances of protist functional groups observed in our experimental conditions (flagellates > amoebae > ciliates) matched *in situ* observations from soil protist communities [19, 20]. Collectively, these results indicate that microbial assemblages developing in the chips accurately reflected the taxonomic and functional diversity, and trophic structures of microbial communities colonizing organic matter patches in soil.

Based on microscopic counts, Hypotrichia individuals represented only a very small fraction— approximately 0.18%—of the protist community in control chips. Their low abundance and consistent presence made them suitable candidates to assess as potential keystone species [3]. Although microscopy suggested a single Hypotrichia morphotype/species was present, DNA metabarcoding indicated the likely presence of three distinct Hypotrichia species, with being one overwhelmingly more abundant and prevalent than the others. While these latter results suggest that technically we manipulated the abundance of keystone functional guild [10], the notably higher abundance of OTU30 relative to the other Hypotrichia OTUs indicates our suppression was largely of a single keystone species. We also recognize that our Hypotrichia suppression was a full removal treatment, due to rapid recolonization from surrounding soil. Sealing the chip entrance post-inoculation following ciliate colonization (around day 8 in our experimental conditions) in combination with selective suppression might have achieved full suppression; however, this approach would reduce ecological relevance, as microbial communities rapidly succeed during organic matter decomposition, and our chip design mimicked recently senesced fungal necromass [51].

Contrary to our first hypothesis—that Hypotrichia suppression would decrease bacterial diversity by releasing predation-resistant taxa to dominate—we found bacterial diversity and community composition remained remarkably stable across treatments. We suspect this unexpected stability likely resulted from the compensatory increase in flagellate abundance observed following Hypotrichia suppression. Flagellates, known bacterial predators [52–54], likely occupied the predatory niche vacated by Hypotrichia, maintaining bacterial community stability. Alternatively, it is possible that the bacterial communities in our experimental conditions may have inherently resisted protist predation, remaining unaffected despite increased flagellate abundance. Consistent with that possibility, we observed bacterial biofilm formation in the chips, which may have limited protist predation. Furthermore, other studies has shown limited top-down control of bacterial populations by ciliates or flagellates [20, 55, 56], with Clarholm in 1981 specifically highlighting amoebae as more effective bacterial predators in soil systems [57]. Given that preliminary experiments indicated our phototoxicity application could also effectively suppress large amoebae, targeting amoebae in future keystone species experiments would be valuable.

Partially supporting our second hypothesis, Hypotrichia suppression significantly decreased fungal diversity, with fungal abundance and richness also marginally declining. Initially, we expected fungal community changes resulting from altered bacterial competition. However, bacterial abundance, richness, and composition were unchanged, suggesting other mechanisms drove fungal decline. We postulate that decline was likely due to increased flagellate abundance. The dominant flagellates (e.g., *Cercomonas*) present on our chips are facultative fungal predators[17], suggesting that Hypotrichia suppression indirectly intensified flagellate top-down control on fungi. This change in fungal diversity, most likely mediated by non-target predator abundance alterations, further aligns with the keystone species concept.

Supporting our third hypothesis, we observed clear shifts in protist community abundance and richness following Hypotrichia suppression, prominently marked by increased flagellate abundance. These findings strongly parallel the mesopredator release concept in animal ecology, where removing apex predators leads to subordinate predator proliferation [58, 59]. The significant negative correlation between flagellates and suppressed Hypotrichia suggests competition for bacterial prey; however, direct Hypotrichia predation on flagellates cannot be excluded, as some Hypotrichia may consume small flagellates [26, 60]. Regardless of the specific mechanism, our results align with studies reporting negative interactions—through competition or predation—between ciliates and flagellates, particularly involving ciliates larger than our experimental Hypotrichia [61–63]. Overall, we observed that Hypotrichia suppression significantly increased protist diversity by enabling flagellate proliferation from diverse lineages (e.g., Bigyra, Choanoflagellata, Euglenozoa) that were previously rare or absent on control chips.

Further support for our third hypothesis emerged from protist community compositional shifts, with Hypotrichia suppression resulting in more similar communities. This result represents a microscale example of biotic homogenization [64, 65] and also Paine’s findings in the original studes of the keystone concept [1, 2]. Although biotic homogenization here coincided with increased protist community diversity, these processes are not mutually exclusive, leaving open questions about potential broader cascading effects on community stability and function over time. For example, unexpectedly, Hypotrichia suppression correlated with reduced non-Hypotrichia ciliates (*Platyophrya*, *Microthoracida*), suggesting possible facilitative interactions among ciliates, although the mechanisms remain unclear. Taken together, these observations closely align with the original conceptualization of keystone species as significantly affecting community structure irrespective of whether its effects were beneficial or detrimental [1, 2].

While our study provides robust evidence that suppressing a low abundance microbial predator species can induce major changes in microbial community composition and diversity, some limitations must be acknowledged. First, we observed surprisingly high within-treatment variability for microbial metrics despite using the same homogenized soil pool as inoculum. We suspect this variability is likely linked to priority effects, where the identity of the first chip microbial colonizers (bacteria, fungi, or protists) influenced subsequent community assembly [66]. Such variability reduced the power of our statistical analyses, especially when a few chips exhibited unusual patterns—such as extremely low total protist counts or an early peak followed by a rapid decline in Hypotrichia in the control treatment. As such, establishing pre-determined exclusion criteria for outlier chips could help mitigate these issues in future experiments. Another limitation was the presence of nematodes, which, although not significantly different between treatments and primarily inactive or dead within the chips, dominated the 18S sequencing dataset and reduced protist community read depth. Developing a chip entrance system to exclude nematodes may improve the quality of protist sequencing data. It is also important to note that our endpoint experiment did not capture the longer-term temporal dynamics of microbial communities and microbial community stability—a key component of the keystone species concept [1–3]—so incorporating time-series sampling in future studies will be particularly important. Finally, while phototoxicity proved effective for suppressing large protists like ciliates, future developments could involve AI-driven training of protist species morphologies combined with an automated high-magnification suppression system (e.g., 1000×), enabling highly efficient, species-specific, and temporally continuous suppression protist species within synthetic soil spore space systems.

## Conclusions

Here, we deployed a novel experimental system to directly test the keystone species concept within microbial communities by integrating microfluidic soil chips—allowing complex microbial communities to develop under semi-controlled conditions—with the selective suppression of a protist predator through targeted phototoxicity. To our knowledge, this study provides the first experimental evidence demonstrating that a protist predator can function as a keystone species within a complex soil microbial community. Specifically, our findings validate key tenets of the keystone species concept, illustrating that a low-abundance predator species can disproportionately influence community diversity and composition. Future research prioritizing time-series analyses and investigating the functional consequences of suppressing keystone species will help to further clarify the role of microbial predators in regulating community stability and influencing soil ecosystem processes over extended periods.

## Supplementary material

**Table S1.** Summary of chips used across experiments, detailing treatments and reasons for exclusion.

**Table S2.** Taxonomic assignments for protist functional groups.

**Figure S1.** (a) Design schematic of the chips used in the study, comprising a 14,720 µm × 5000 µm cuboid structure with a pillar network (175 µm center-to-center spacing); (b) Representative microscopy image of a chip bonded to a glass coverslip, inoculated with fungal necromass and soil.

**Figure S2.** Representative microscopy images illustrating bacterial abundance scoring in chips, ranging from 1 (very low abundance) to 5 (very high abundance). Scale bar = 40 µm.

**Figure S3.** Microscopy image demonstrating bacterial counting by deep-learning algorithm (Zou et al., 2024). Light blue squares indicate counted bacterial cells; the dark red square indicates a bacterial cluster excluded from the analysis. Scale bar = 40 µm.

**Figure S4.** Representative microscopy images for counting protist functional groups: (a) Hypotrichia (suppressed), (b) non-Hypotrichia ciliates, (c) amoebae, (d) flagellates, and (e) fungal hyphae. Scale bar = 25 µm.

**Figure S5.** Hypotrichia abundance and identification: (a) Relative abundance in protist metabarcoding dataset (dark dots are means ± SE, lighter dots represent individual chips, statistical comparisons via t-test); (b) 18S sequence similarity matrix among six Hypotrichia-assigned OTUs; (c) Neighbor-Joining phylogenetic tree showing sequence similarity, relative abundances (relative to total Hypotrichia), and frequencies across chips.

**Figure S6.** Relative abundance of nematodes and rotifers (compared to total protist OTUs) by treatment (control vs. Hypotrichia suppression); dark dots show means ± SE, lighter dots individual chips. Nematode abundances compared by t-test; rotifer abundance not statistically analyzed due to single-sample presence.

**Figure S7.** Effects of Hypotrichia suppression on bacterial abundance from microscopy scoring (1– 5 scale, see Figure S2). Dark dots represent means ± SE; lighter dots individual chips. Statistical comparisons via t-tests or Welch’s tests.

**Figure S8.** Relative abundance of dominant bacterial genera by treatment (control vs. Hypotrichia suppression).

**Figure S9.** Relative abundance of dominant fungal genera by treatment (control vs. Hypotrichia suppression).

**Figure S10.** Differences in relative abundances of protist OTUs between control and suppressed treatments. OTUs summed at genus or higher taxonomic level when genus-level data was unavailable; numbers in brackets indicate overall relative abundance in the metabarcoding dataset.

**Figure S11.** Pearson’s correlations between protist functional groups and Hypotrichia abundance (day 20): (a) flagellate abundance (microscopy), (b) flagellate relative abundance (metabarcoding), (c) amoeba abundance (microscopy), (d) amoeba relative abundance (metabarcoding), (e) ciliate abundance (microscopy), (f) ciliate relative abundance (metabarcoding). Significance marked by asterisks (\**p* < 0.05, \*\**p* < 0.01, \*\*\**p* < 0.001).

## Author’s contributions

Conceptualization: F.M.

Data curation: F.M., F.K., H.Z.

Formal analysis: F.M., F.K., H.Z.

Funding acquisition: F.M., A.T., E.C.H., P.G.K.

Investigation: F.M., F.K., B.H.B.

Methodology: F.M., H.Z., E.C.H.

Project administration: F.M.

Resources: F.M., A.T., E.C.H., P.G.K

Software: H.Z.

Supervision: F.M.

Visualization: F.M., F.K.

Writing – original draft: F.M.

Writing – review & editing: all authors

## Acknowledgements

The authors acknowledge financial support provided by the Swedish Research Council (Starting grant within natural and engineering sciences, VR 2023-04643) and the Crafoord Foundation (Research grant, Crafoord 20241084) to F. Maillard, a Research Grant in Natural and Engineering Sciences to A. Tunlid (VR 2021-05188), the Wardle Chair of Microbial Ecology Fund to P.G. Kennedy, and a Future Research Leader grant (FFL18-0089) by the Foundation for Strategic Research to E.C. Hammer. We acknowledge the strategic research environment Biodiversity and Ecosystem Services in a Changing Climate, BECC, funded by the Swedish Government. We thank P.M. Mafla-Endara for her help with the microscopy setup and the University of Minnesota Genomics Center (UMGC) for their support in concentrating and sequencing the HTS libraries.

