## Supplemental Methods for "Experimental suppression of a keystone protist triggers mesopredator release and biotic homogenization in complex soil microbial communities"

**Supplementary Materials and Methods**

*Chip design and fabrication*

The microfluidic chip design consisted of a cuboid space measuring 14,720 μm in length, 500 μm in breadth, and 12 μm in height, within which circular pillars approximately 100 μm in diameter were arranged in a regular array with a center-to-center spacing of 175 μm (see Figure S1a). This relatively small chip format was chosen to allow rapid inspection at low magnification (40×) and to facilitate the identification and suppression of relatively large protists. Chips were fabricated by molding PDMS (Sylgard 184, Dow Corning) on a photoresist master produced via UV lithography with SU-8 5 (MicroChem Corp), following the protocols of Aleklett et al. (2021) and Mafla-Endara et al. (2021). To produce the master, a thick negative photoresist (SU-8 5, MicroChem Corp, USA) was spin-coated onto a glass plate at 1250 rpm for 60 s. This layer was soft baked on a hot plate at 90 °C for 5 min, patterned using UV exposure with a Karl-Suss MA4 mask aligner, and post-exposure baked. The photoresist was then developed for 3 min in mr-Dev 600 (MicroChem) and rinsed with isopropanol (VWR International). PDMS slabs were produced by mixing the PDMS base and curing agent (both Sylgard 184, Dow Corning, USA) in a 10:1 mass/mass ratio. The mixture was poured onto the master in a 4-mm-thick layer and degassed in a vacuum chamber at −25 kPa for 45 min. The PDMS was cured in an oven at 60 °C for 2.5 h. Once cooled, the PDMS was cut slightly larger than the patterned area, and an entrance was created by excising one long side of the PDMS slab. Finally, the PDMS slabs were bonded to glass slides. Glass slides (55 × 75 mm; Thermo Scientific) were cleaned sequentially with acetone and 75% ethanol. Both the PDMS and glass slides were treated separately in an oxygen plasma chamber (Diener Electronic Zepto): the glass slide for 1 min under UV light and the PDMS for 10 s. Immediately after treatment, the activated surfaces were brought together and gently pressed in the central region of the chip.

*eDNA extraction*

After video recording at day 20, any soil on each chip was carefully removed. The chip contact area with soil was cleaned multiple times using a cotton swab moistened with molecular-grade water followed by a dry cotton swab to remove residual soil or necromass particles, thereby minimizing soil eDNA contamination. Using tweezers, the chip was detached from the coverslip, flipped over, and transferred to a Petri dish. The glass coverslip area that had been in contact with the chip was swabbed with a moistened cotton swab, and the tip was immediately placed into the extraction tube of the PowerSoil Pro Kit (Qiagen, Hilden, Germany). Using a sterile scalpel blade, the edges of the chip were excised to remove potential surface contaminants, and the remaining portion—representing the imprint of the pillar array in the PDMS—was transferred directly into the extraction tube. This procedure ensured that microbial cells adhering to both the coverslip and PDMS were collected. Genomic DNA was then extracted according to the PowerSoil Pro Kit manufacturer’s protocol, with a final elution volume of 50 μL. Three non-inoculated chips were processed identically as negative controls.

*Microbial qPCR*

Bacterial abundance was quantified via qPCR on a Stratagene Mx3005P PCR machine (Agilent Technologies), targeting the bacterial 16S rRNA gene with the 1401F/968R primer set (Cébron et al., 2008). Each 20 μL qPCR reaction contained 4 μL of template DNA, a standard 16S bacterial double-stranded DNA template (ranging from 10⁹ to 10³ gene copies μL⁻¹) or molecular-grade water as a negative control, and SsoAdvanced Universal Inhibitor-Tolerant Supermix (Bio-Rad, Hercules, CA, USA). The amplification protocol included an initial denaturation at 95 °C for 5 min, followed by 40 cycles of 20 s at 95 °C, 30 s at 56 °C, and 60 s at 72 °C. Primer specificity was confirmed via melting curve analysis from 70 °C to 95 °C, increasing at 0.3 °C s⁻¹. All samples were run in technical duplicates, and results were averaged to obtain the bacterial 16S copy number per chip. An attempt to quantify fungal abundance using qPCR with the FR1/FF390 primer set (Chemidlin Prévost-Bouré et al., 2011) was unsuccessful, as all samples fell outside the standard curve very likely due to low fungal genomic DNA concentrations.

*High-throughput amplicon sequencing*

To characterize microbial community structure within the chips, high-throughput sequencing (HTS) of taxonomic markers for bacteria, fungi, and protists was performed. Sixteen chip samples (from the 20 initially inoculated, with four excluded as previously noted) and three negative control chips were analyzed. Bacterial community structure was assessed by amplifying the V4 region of the 16S rRNA gene using the 515F–806R primers (Caporaso et al., 2012). Fungal community structure was characterized by targeting the ITS2 region with the 5.8S-Fun and ITS4-Fun primers (Taylor et al., 2017). For evaluating protist community structure, the 18S rRNA gene was amplified using the 616*f–1132r primer pair (Hugerth et al., 2014), which captures a broad taxonomic range of eukaryotes (Vaulot et al., 2022). First-round PCRs (20 μL reactions) contained 10 μL of Phusion Hot Start II High-Fidelity PCR Master Mix (Thermo Scientific, Waltham, MA, USA), 0.5 μL of each 20 mM primer, and 10 μL of template DNA. Preliminary tests indicated that replacing the water in the PCR mix entirely with template DNA improved amplification from the limited number of microbial cells in the chips. The thermocycling conditions were as follows: an initial denaturation at 98 °C for 30 s; then 39 cycles of 98 °C for 30 s, annealing at 50 °C (16S), 55 °C (ITS), or 52 °C (18S) for 30 s, and extension at 72 °C for 30 s; with a final extension at 72 °C for 10 min. A second PCR was performed to add unique Golay barcodes and sequencing adapters. PCR products were purified and normalized using the Charm Just-a-Plate Purification and Normalization Kit (Charm Biotech, San Diego, CA, USA) and pooled at equimolar concentrations. Sequencing was conducted on an Illumina MiSeq platform using 2 × 300 bp chemistry at the University of Minnesota Genomics Center.

Sequencing data for 16S, ITS, and 18S amplicons were processed using the DADA2 pipeline in R (Callahan et al., 2016). For the 18S and ITS datasets, only forward reads (R1) were used to mitigate data loss due to poor R2 quality in combination with longer amplicon that requires relative high-quality score for efficient paired-end merging; paired-end merging was used for the 16S dataset. Raw fastq files were filtered to remove reads with ambiguous bases (maxN = 0), and primers were removed using Cutadapt (Martin, 2011). Quality filtering was applied with the following parameters: for 18S, a truncation length of 210 bp and maxEE = 25; for ITS, truncLen = 210 bp and maxEE = 8; and for 16S, forward and reverse reads were truncated at 230 and 150 bp with maxEE values of 8 and 12, respectively. Error rates were learned, and amplicon sequence variants (ASVs) were inferred using DADA2, with chimeric sequences removed using removeBimeraDenovo. Taxonomy was assigned using the assignTaxonomy function with the PR2 database for 18S (Vaulot et al., 2022), the UNITE database for ITS (Abarenkov et al., 2024), and the SILVA database for 16S (Quast et al., 2012). ASV sequences were then aligned using DECIPHER’s AlignSeqs, a distance matrix computed, and sequences clustered into operational taxonomic units (OTUs) at a 97% similarity threshold using DECIPHER’s TreeLine.

Following OTU clustering, OTU tables were run through additional quality control. OTUs detected in the three non-inoculated control chips were removed from the chip samples (11 OTUs for bacteria, 3 for fungi, and 3 for protists). Additionally, OTUs lacking kingdom-level taxonomic information were blasted against NCBI to confirm affiliation with the expected kingdom (bacteria, fungi, or protists), and those failing to match were omitted (8 OTUs for bacteria, 11 for fungi, and 3 for protists). OTUs corresponding to organelles (e.g., chloroplasts, mitochondria) were also removed (9 OTUs from the bacterial dataset). For the protist dataset, further filtering removed 72 bacterial OTUs, 35 fungal OTUs, 37 metazoan OTUs, and 1 plant (*Picea*) OTU. Metazoan OTUs were subsequently subset between nematodes and other microscopic animals (in this case, rotifers) to assess their potential impact on the results. The bacterial and fungal OTU tables were normalized by rarefaction using “rrarefy function” in R Vegan package (Oksanen et al., 2015) to 1716 and 2578 reads, respectively; two chips were removed from the bacterial analysis due to low sequence counts (533 and 624 reads) and three chips were excluded from the fungal analysis (with 0, 1, and 5 reads). To examine the effects of Hypotrichia suppression on protist community composition and diversity, all Hypotrichia sequences were removed from the protist dataset before the analyses. Due to the overall low read depth (average ~1060 sequences), normalization of this dataset was performed using relative abundances rather than rarefaction. One chip was further omitted because it yielded only a single 18S OTU, despite microscopy indicating a more diverse protist community. A summary of the chips retained for the community analyses is provided in Table S1. Finally, to facilitate direct comparison between microscopy and metabarcoding results, protist OTUs were regrouped into functional groups based on taxonomic information (see Table S2).
